# Physiological, Histological, and Cognitive Characterization of a Rhesus Macaque Model of Presbycusis

**DOI:** 10.64898/2026.03.24.714040

**Authors:** Swarat Kulkarni, Amy Conner, Oscar Rausis, David Pitchford, Zhengyang Wang, Aneesh Batchu, Leslie Liberman, M. Charles Liberman, Christos Constantinidis, Troy Hackett, Ramnarayan Ramachandran

## Abstract

Age-related hearing loss (ARHL), or presbycusis, is one of the most prevalent sensory deficits in older adults and has been increasingly implicated in cognitive decline and dementia. This study characterizes ARHL in a rhesus macaque model by combining histological, physiological, and cognitive assessments. Aged macaques exhibited progressive cochlear degeneration, with marked outer hair cell loss at mid-to-high frequencies, elevated auditory thresholds, reduced distortion product otoacoustic emissions, and impaired auditory brainstem responses including amplitude reduction, latency prolongation, and diminished temporal precision. Despite modest reductions in inner hair cell ribbon synapse counts, hypertrophic changes were observed. These auditory deficits correlated with subtle impairments in visual working memory, as measured by a delayed match-to-sample task, underscoring a potential sensory-cognitive link. By capturing cross-domain aging markers in a translationally relevant primate model, this work lays a foundation for mechanistic studies and therapeutic interventions targeting both hearing and cognition in aging populations.

## Introduction

Age-related hearing loss (ARHL), also known as presbycusis, affects nearly two-thirds of the U.S. population aged 70 and older (Tawfik et al., 2019). Although it is common, ARHL is not a benign consequence of aging. It is associated with increased healthcare utilization due to its links with falls, depression, social isolation, cognitive decline, and dementia (Sharma et al., 2021; Choi et al., 2016), contributing to an estimated $3 billion in annual medical expenditures (Foley et al., 2014). As the aging population grows, the number of individuals affected by ARHL and the associated economic burden is expected to rise substantially (Lin et al., 2013). Although hearing aids and cochlear implants can restore audibility and improve communication (Choi et al., 2016; Contrera et al., 2016), fewer than 4% of adults who could benefit from amplification use it (Chien, 2012; Lin et al., 2011). This disconnection between high prevalence, low treatment uptake, and unclear benefit highlights the need for a deeper mechanistic understanding of ARHL and its broader effects on brain and behavior.

Although age-related hearing loss is often characterized as a peripheral sensory disorder, its broader impact on auditory processing and cognition is increasingly recognized. Peripheral decline, particularly when affecting signal fidelity and temporal precision, places additional demands on central auditory pathways and increase the cognitive effort required for everyday listening (Bush et al., 2015). Over time, this increased listening effort has been hypothesized to divert cognitive resources away from other domains such as memory and executive function. Communication difficulties may also contribute to social withdrawal and reduced cognitive stimulation, both of which are associated with elevated risk for cognitive decline (Samtani et al., 2022). Longitudinal studies have shown that individuals with untreated hearing loss have a 30–40% accelerated rate of cognitive decline and a 24% increased risk of incident cognitive impairment over a 6-year period compared to age-matched peers with normal hearing (Lin et al., 2013). These findings underscore the importance of understanding how structural changes in the peripheral auditory system may relate to functional auditory deficits and early cognitive decline.

Rodent and human studies have provided substantial insight into the cellular basis of auditory aging. Histological analyses reveal progressive loss of outer and inner hair cells (OHCs and IHCs), often accompanied by afferent synapse loss and degeneration of spiral ganglion neurons (Spongr et al., 1997; Wu et al., 2019; Ueno et al., 2026). In rodents, OHC loss tends to occur earliest in the basal and apical regions, producing a U-shaped pattern of vulnerability across the cochlear length, while mid-frequency regions are relatively preserved (Spongr et al., 1997). In humans, postmortem analyses show comparable but more variable patterns of degeneration, with extensive OHC loss, moderate IHC loss, and pronounced degeneration of the auditory nerve that often exceeds the degree of IHC loss (Wu et al., 2019). This dissociation suggests that cochlear synaptic integrity may decline independently from sensory cell degeneration, and that cumulative lifetime exposures—such as noise, ototoxic medications, or age-related vascular insufficiency affecting the stria vascularis—may accelerate synapse loss even in individuals with clinically “normal” thresholds (Valderrama et al., 2018; Sergeyenko et al., 2013). Physiologically, both rodents and humans demonstrate age-related declines in cochlear amplification, reduced distortion product otoacoustic emission (DPOAE) amplitudes, elevated auditory brainstem response (ABR) thresholds, and delayed neural latencies (Abdala & Visser-Dumont, 2001; Konrad-Martin et al., 2012).

Despite these similarities, important translational gaps remain. Rodents experience accelerated cochlear aging, shorter lifespans, and heightened susceptibility to acoustic trauma (Gray & Barnes, 2019). Human studies lack experimental control and are confounded by decades of variable environmental, vascular, and medical exposure (Vaden et al., 2022; Valderrama et al., 2018). These limitations highlight the need for an intermediate model that bridges the experimental precision of rodents with the anatomical and behavioral complexity of humans. Nonhuman primates, such as rhesus macaques, offer such a model: they are relatively resistant to acoustic trauma and share key structural and functional features with humans, making them a promising yet incompletely characterized system for studying age-related hearing loss (Chiou et al., 2020; McLeod et al., 2022). Importantly, macaques can be assessed across molecular, physiological, and behavioral domains, enabling investigation of how peripheral degeneration relates to functional hearing loss—defined as measurable deficits in auditory processing (e.g., degraded temporal precision, reduced amplitude of auditory brainstem responses, or impaired sound discrimination) that manifest even when other aspects of audition are preserved.

Here, we leveraged the rhesus macaque model to examine the relationship between cochlear degeneration, auditory processing deficits, and early signs of cognitive decline. We combined histological analysis of hair cell survival and afferent synapse integrity with physiological assessments of cochlear and brainstem function using DPOAEs and ABRs, as well as behavioral memory testing. Our goal was to elucidate the relationship between structural changes in the cochlea, functional hearing loss, and emerging cognitive vulnerability in a translationally relevant primate model, providing a foundation for future investigations into the mechanisms and progression of auditory and cognitive aging. We hypothesized that age-related loss of cochlear integrity would be associated with measurable declines in auditory temporal processing and brainstem synchrony, which in turn would predict subtle impairments in cognitive performance.

## Methods

### Subjects

Nine aged macaque monkeys (Macaca mulatta; n = 6 males, n = 3 females; ages 26–34 years) were included in this study. One additional animal received auditory testing but was ultimately excluded from analysis due to limited testing capacity and health complications. These animals, referred to in this manuscript as old macaques, were previously a part of the Pennington et al. study (2025). For comparison, previously published data from 18 young macaques (*Macaca mulatta*, n=34 ears, and *Macaca radiata*, n=2 ears; ages 6-10 y/o) were included from Stahl et al. (2022).

Subjects were further categorized by age into young-old (26–30 years, n = 12 ears) and old-old (31–34 years, n = 6 ears). These age ranges span the upper end of the macaque lifespan in captivity, which has been reported to exceed 30 years (Chiou et al., 2020; Roth et al., 2004). Based on established lifespan conversions, one macaque year approximates three human years (Davis and Leathers, 1985), making the OM cohort roughly equivalent to 78–102 human years. Auditory data were collected shortly before planned euthanasia, and cochlear tissues were preserved for histological analysis with approval from the Constantinidis Lab. Five of the nine old monkeys had previously participated in a separate study in the Constantinidis Lab involving electrical stimulation to evaluate effects on cognitive function during aging (Pennington et al., 2025).

All housing and procedural protocols were approved by the Institutional Animal Care and Use Committee (IACUC) at the Vanderbilt University Medical Center and were in strict compliance with the guidelines established by the National Institutes of Health (NIH). Subjects were excluded from the study if they possessed abnormal outer or middle ear function as measured during otoscopy and tympanometry.

All macaques used in this study were housed at Vanderbilt University on a 12-hour light/dark cycle and provided access to a carefully controlled diet *ad libitum*, except for 12–18 hours prior to sedated physiological testing. Animals were socially paired when possible to promote social enrichment. All subjects had access to a range of environmental enrichment materials, including foraging devices, and were housed in rooms containing multiple animals. Monkeys were maintained in typical colony acoustic environments with variable background noise from vocalizations and socialization activities.

### Sedation

Monkeys were fasted for 12-18 hours before the sedated physiological testing procedure. Subjects were anesthetized with a cocktail of ketamine (10-13 mg/kg IM) and midazolam (0.1-0.3 mg/kg IM) and treated with glycopyrrolate (0.004-0.015 mg/kg) to minimize mucous secretions. They were then intubated and transferred to a sound-treated room (Acoustic Systems ER247), switched to a breathing circuit, and maintained with 1-2% isoflurane for the remainder of testing.

### Otoscopy

Visual inspection of the external auditory canal and tympanic membrane was performed using an otoscope (Welch Allyn, 25020 3.5V Halogen Otoscope Head) to assess for contraindications to completing physiological testing. These contraindications included excessive cerumen and/or debris, active otorrhea, and tympanic membrane perforation, bulging, or retraction. All macaques included in this study had minimal to no non-occluding cerumen.

### Tympanometry

Following otoscopic inspection, standard clinical tympanometry was completed using a calibrated Amplivox Otowave tympanometer. A 226-Hz probe tone was generated in the ear canal while the ear canal pressure was systematically swept from +200 to −300/400 daPa. Changes in probe level during the pressure sweep resulted in estimates of tympanic membrane compliance (mL), middle ear pressure (daPa), and ear canal volume (ECV, cc). Subjects with very large ear canal volumes (>1.0 cc), very small ear canal volumes (<0.2 cc), or middle ear pressure values that were < −150 daPa were excluded from the study due to likely tympanic membrane perforation, cerumen impaction, or middle ear dysfunction. Accordingly, all rhesus macaque subjects met the inclusion criteria and proceeded to subsequent physiological, histological, and cognitive testing. Tympanometry from young-old and old-old macaques did not significantly differ in compliance, pressure, or ECV (p > 0.05, Welch’s ANOVA and Games-Howell post-hoc) from normative data collected from macaques (Stahl et al., 2022), and all individual monkey values were considered normal.

**Table 1:** Mean plus or minus one Standard Error of the Mean (SEM) of tympanometric measures. *Note*: cc = cubic centimeters, mmho = millimho, daPa = decapascals.

| Age Group | ECV (cc) | Pressure (daPa) | Compliance (mL) |
| --- | --- | --- | --- |
| Young (6-10 y/o) | $0.400 \pm 0.031$ | $-9.00 \pm 5.28$ | $0.50 \pm 0.03$ |
| Young-old (26-30 y/o) | $0.550 \pm 0.041$ | $2.00 \pm 2.15$ | $0.55 \pm 0.06$ |
| Old-old (31-34 y/o) | $0.433 \pm 0.047$ | $3.17 \pm 19.5$ | $0.45 \pm 0.11$ |

### Distortion Product Otoacoustic Emissions (DPOAEs)

A high frequency DPOAE system (ER-10X, Etymotic) was used to measure DPOAEs at 4 frequencies per octave from *f*_2_ = 1-32 kHz. For each ear, distortion product otoacoustic emission (DPOAE) amplitude (signal level) and the noise floor were measured across all stimulus level and frequency combinations. From these measurements, DP-grams were generated, which plot DPOAE thresholds or amplitudes across each tested DP frequency. Thresholds were defined as the lowest stimulus level at which a distortion product was considered present (the DP amplitude exceeded 0 dB SPL and was at least 6 dB above the noise floor), following criteria used by the Vanderbilt Audiology Clinic. DP-grams were measured in all subjects using an *L*_1_/*L*_2_ ratio of 65/55 and an *f*_2_/*f*_1_ ratio of 1.22. The DPOAEs collected over a range of stimulus levels with *L*_1_ – *L*_2_ = 10 dB were used to derive threshold-versus-age functions at each frequency. These frequency and level ratios of the primary tones were developed from previous study of young monkeys (Stahl et al., 2022).

### DPOAE Analysis

Threshold was estimated as the lowest f_2_ level that produced a response meeting these presence criteria. Because the step size between stimulus levels was 5 dB, this often resulted in multiple ears sharing the same threshold value. To improve precision, SNR versus f_2_ level functions were fitted with a psychometric curve, and threshold was estimated from the fitted function.

### Auditory Brainstem Responses (ABRs)

Auditory brainstem responses (ABRs) were recorded using a vertex-to-mastoid electrode montage in response to three types of acoustic stimuli: broadband clicks (0.001–97 kHz; 100 µs duration; 27.7/s), chirps (0.5–32 kHz; 1.6 ms duration; 27.7/s), and tone bursts (0.5–32 kHz; 27.7/s) spanning much of the audible range of macaques (Pfingst et al., 1978). Additional ABRs were collected in response to clicks presented at varying rates (27.7, 57.7, 100, 125, 166.6, and 200 clicks/s; 70–90 dB SPL).

The recordings used subdermal needle electrodes (Rhythmlink) for the active (mastoid), reference (vertex), and ground (shoulder) placements. Electrode impedances were maintained at ≤ 3 kΩ. A calibrated closed-field speaker (MF1, Tucker-Davis Technologies) was then coupled to a disposable foam eartip (ER3-14 B 10mm, Etymotic Research) to monaurally present stimuli during recordings. Closed-field stimuli were calibrated (+/- 3 dB) using a 0.5cc coupler and verified in the ear canal using a probe microphone system (Fonix 8000, Frye). Stimuli were created in SigGenRZ software (Tucker-Davis Technologies), generated by a RZ6 Multi-I/O Processor (Tucker-Davis Technologies) using BioSigRZ software (Tucker-Davis Technologies), and presented using an alternating stimulus polarity for two separate runs of 1024 presentations. Recordings were made using a low-noise Medusa4Z pre-amplifier with a unity gain setting.

Tone burst stimuli had frequency-dependent rise/fall times and plateau durations consistent with parameters used by the Vanderbilt Audiology Clinic. Specifically, rise/fall times were 1 ms for 2–32 kHz, 2 ms for 1 kHz, and 4 ms for 0.5 kHz; plateau durations were 0.5 ms (2–32 kHz), 1 ms (1 kHz), and 2 ms (0.5 kHz).

### ABR Data Analysis

ABR analysis was performed using the methods described in Stahl et al. (2022). Briefly, ABR waveforms were visually inspected, and peak-to-trough amplitudes and peak latencies were measured for each identifiable wave. Latencies were defined as the time of wave onset (peak), and ABR *threshold* was the lowest stimulus level at which a repeatable, visually detectable wave was present. Chirp analysis followed the same procedures as click analysis, with one exception: all chirp latency values were corrected by subtracting 1.6 ms to account for the stimulus duration inherent to the chirp design. Statistical data processing and analysis were performed using Python, R Studio, and MATLAB. Visualizations were created using Figure Composer.

### Histology

Upon the completion of physiological data collection, subjects (10 young-old ears from 7 animals, 4 old-old ears from 3 animals, and 10 young macaque ears from 7 animals) were euthanized via intravenous overdose of sodium pentobarbital and sodium phenytoin (Euthasol; >120 mg/kg. Cochlear tissue, with surrounding temporal bone, was extracted and perfused through the scala tympani then submerged in 4% phosphate-buffered paraformaldehyde (PFA) for two hours before transfer to 0.12 M EDTA for decalcification. Following decalcification, cochleae were dissected, imaged, and processed for immunohistochemical analysis following established methodology (Valero et al., 2017; Burton et al., 2020; Mondul et al., 2026).

Each cochlear spiral was separated into 10 pieces to isolate whole-mount epithelial preparations of the organ of Corti, encompassing the hair cells and osseous spiral lamina from base to apex. Immunolabeling was performed to visualize: (i) presynaptic ribbons (mouse IgG1 anti-CtBP2; BD Transduction Labs; 1:200), (ii) glutamate receptor patches (mouse IgG2 anti-GluA2; Millipore; 1:200), and (iii) hair cell cytoplasm (rabbit anti-myo7a; Proteus Biosciences; 1:200). A fourth channel was used to label both cochlear efferent terminals (goat anti-choline acetyltransferase (ChAT); Millipore #AB144P; 1:100) as well as stereocilia bundles (rabbit anti-espin; Sigma, #HPA028674; 1:200). Tissue was incubated in appropriate secondary antibody conjugates coupled with AlexaFluors or Pacific Blue fluorophores.

### Histology Analysis

Tissue samples were imaged using a Leica SP8 confocal microscope with a 63X glycerol-immersion objective (1.3 N.A.). Confocal z-stacks were acquired at half-octave intervals along the cochlear spiral, covering frequencies from 0.125 to 32 kHz. Frequency assignments for each stack were derived from a cochlear frequency map (Greenwood, 1990), with an assumed upper frequency limit of 45 kHz.

Hair cell survival was assessed from low-power confocal z-stacks (1024 x 1024 pixels at 141 nm /pixel) by counting cuticular plates and normalizing to the expected number of hair cells per row. Afferent synaptic ribbon counts for inner hair cells (IHCs) were quantified from two adjacent high-power confocal z-stacks (1024 x 512 pixels at 75 nm/pixel) from each cochlear location, using Amira software (version 2019.4; Visage Imaging) with the *connected components* function that finds all volumes within each stack for which all pixels are greater than a criterion intensity. Ribbon counts for each region were divided by the number of IHCs in the stack (including fractions), assessed by counting their negatively stained nuclei in the myo7a channel.

### Statistical analysis

Welch’s ANOVA was applied to assess group differences across auditory measures to account for unequal variances and sample sizes among young, young-old, and old-old macaque subjects across frequencies, intensity levels, and stimulation rates. When significant main effects were observed, Games-Howell post-hoc tests were conducted to identify pairwise differences without assuming homogeneity of variance. Violin plot distributions were evaluated using permutation tests, followed by Wilcoxon Signed Rank Tests for post-hoc comparisons between age groups. Scatter plots were analyzed using ordinary least squares regression to test whether slopes differed significantly from zero and to assess model fit. Age group differences in histogram distributions were evaluated using the Kruskal-Wallis test, with Mann-Whitney U tests conducted post-hoc. All statistical tests were selected for their robustness to non-normality and heteroscedasticity, and all analyses were performed with Python, R Studio, and MATLAB.

### Cognitive Assessment

The subjects were tested for their working memory capacity using a delayed-match-to-sample task as part of another study (Pennington et al., 2025). Briefly, monkeys were trained in their home cage to interact with a touch screen and receive appetitive food pellets as rewards. On each trial, the subject pressed a colored square centered on the screen to start and the square disappeared. After a delay, two squares appeared on the left and right side of the screen symmetrically about the midline, one of which was the same color as the square pressed at the beginning of the trial. Pressing the square with matching color resulted in a food reward and the other a timeout. The delay duration was adaptively adjusted where 3 consecutive correct choices caused a 50% increase in delay duration, and an error caused a decrease (“3-up-1-down”). Each session had at most 96 trials with a starting delay of 0.5 seconds.

Three aspects of their cognitive capacity were quantified from task performance, namely working memory threshold, memory reliability, and choice bias. Working memory threshold refers to the memory delay at which the animal would reach asymptotic performance and was quantified by averaging the last 6 delay reversals in each session. For a “3-up-1-down” design, this corresponds to the delay at which the subject would get 79% of the trials correct on average (Levitt, 1971). The memory reliability and choice bias measures were derived from the mixed-effect logistic regression model and respectively captured memory vs. non-memory related aspects of cognition that could impact task performance. The design is specified in equation 1 where: *Y_i_* was the binary response variable at delay *i* for which 1 indicated a response towards a cue on the right side of the screen. *l* was the laterality of the correct cue, with right and left targets dummy coded 1 and −1 respectively. Thus, *β*_0_ was the log odds of choice bias in the baseline period; *β*_1_ was the delay-averaged memory retrieval reliability; and *b_i_* was the random effect of delay *i* on memory retrieval reliability, allowing the estimation of bias across delays.

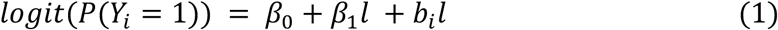

The random-effect coefficients were modeled to follow a normal distribution specified in equation 2. The number of correct and total trials were aggregated across sessions by cue laterality and delay. Only delays at which at least 20 trials were completed across conditions for all subjects were included, ensuring a balanced design for each subject and directly comparable *β*_0_ and *β*_1_ across subjects. *β*_1_ thus measures how reliably the monkey could retrieve the correct choice from memory across delays (reliability), while *β*_0_ measures potential lateralized sensory, motor, or executive deficits that would shift the decision curve (bias), with both represented on a log likelihood scale.

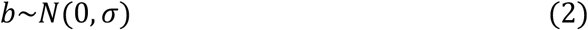

## Results

### Hair cell loss, synaptopathy and stereocilia condition

To investigate age-related cochlear degeneration, we quantified OHC and IHC survival, as well as IHC synaptic ribbon counts, across the cochlear frequency map. As expected, the hair cell loss was most pronounced in the basal half of the cochlea and was more severe for OHCs (Fig. 1E) than for IHCs (Fig. 1F). The inter-animal variability was larger in the oldest animals.

**Figure 1:**
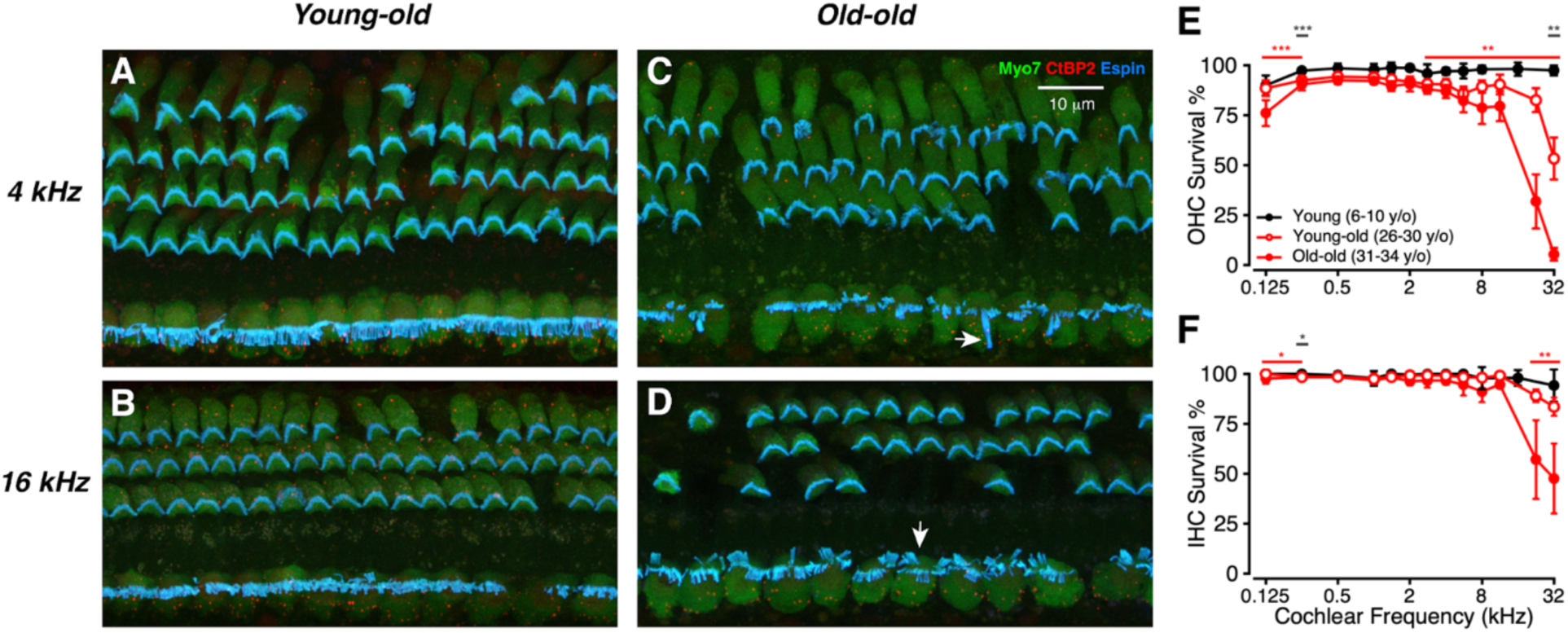
Patterns of hair cell loss in aged macaques. (A-D): Representative confocal projections from the 4 or 16 kHz regions from Young-old or Old-old macaques. Scale bar in C applies to all four images. Arrows in C and D point to abnormal stereocilia bundles on IHCs. (E, F) Mean OHC or IHC Survival (±SEMs) as a function of cochlear frequency for Young (black), young-old (unfilled red, dark gray *), and old-old (filled red, red *) macaques, as indicated in the key used in all following figures unless otherwise noted. Significant p-values are displayed with an asterisk scheme (*: *p* < 0.05, **: *p* < 0.01, ***: *p* < 0.001).

We included an espin stain to allow analysis of stereocilia condition, as seen in the confocal images of Figure 1A-D. Although there were signs of pathology, such as fused, elongated or disarrayed stereocilia on the IHCs in some of the old-old animals, a qualitative analysis suggested no obvious correlation between stereocilia condition and thresholds.

IHC synaptic ribbons were next investigated (Figure 2), with representative young-old and old-old ribbons displayed in Figure 2A-D and average ribbons per IHC summarized in Figure 2E. Synaptic ribbons in the Young-old mice showed the modiolar-pillar gradient in size that has been reported in prior work on guinea pigs and mice (Liberman et al., 2011; Furman et al., 2013), as seen in the zy projections in Figure 2B and D. There was a small but significant reduction in synaptic ribbon counts throughout the cochlear spiral in both the Young-old and Old-old ears, compared with young controls (Fig. 2E). Immunostaining for the GluA2 receptors was much noisier in the Young-old and Old-old ears, compared with the young controls. This made it impossible to reliably distinguish orphan ribbons from those paired with post-synaptic receptor patches. Similar issues have been seen with older mice (Grierson et al., 2022).

**Figure 2:**
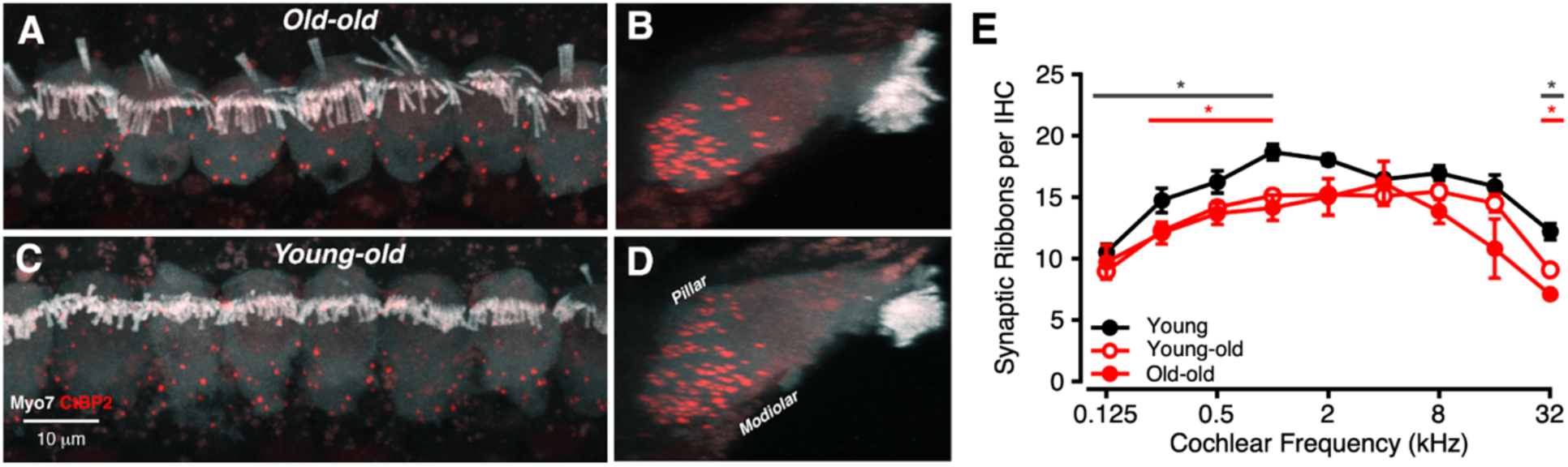
Patterns of synaptic ribbon loss in aged macaques. (A-D): Confocal projections of representative confocal z-stacks from an Old-old and a Young-Old macaque, shown both in the acquisition plane (A, C) and as “side views” (B, D). (E): Mean ribbon counts (±SEMs) for Old-old and Young-old macaques compared to young animals. Significant p-values are displayed with an asterisk scheme (*: *p* < 0.05, **: *p* < 0.01, ***: *p* < 0.001, ****: *p* < 0.0001).

IHC ribbon volumes were also quantified and grouped into frequency bands (Figure 3). In addition to fewer IHC ribbons with age, ribbon volumes vary across frequency bands (low = 0.125-2 kHz, mid = 4-8 kHz, high = 16-32 kHz). At low frequencies (Fig. 3A), young-old macaques had larger ribbon volumes (median = 0.197) than both young (0.183) and old-old (0.181) animals (Kruskal–Wallis with Dunn’s post-hoc, *p* < 0.0001). At high frequencies (Fig. 3C), both young-old (0.227) and old-old (0.232) groups exhibited hypertrophy relative to young macaques (0.210; *p* < 0.0001).

**Figure 3:**
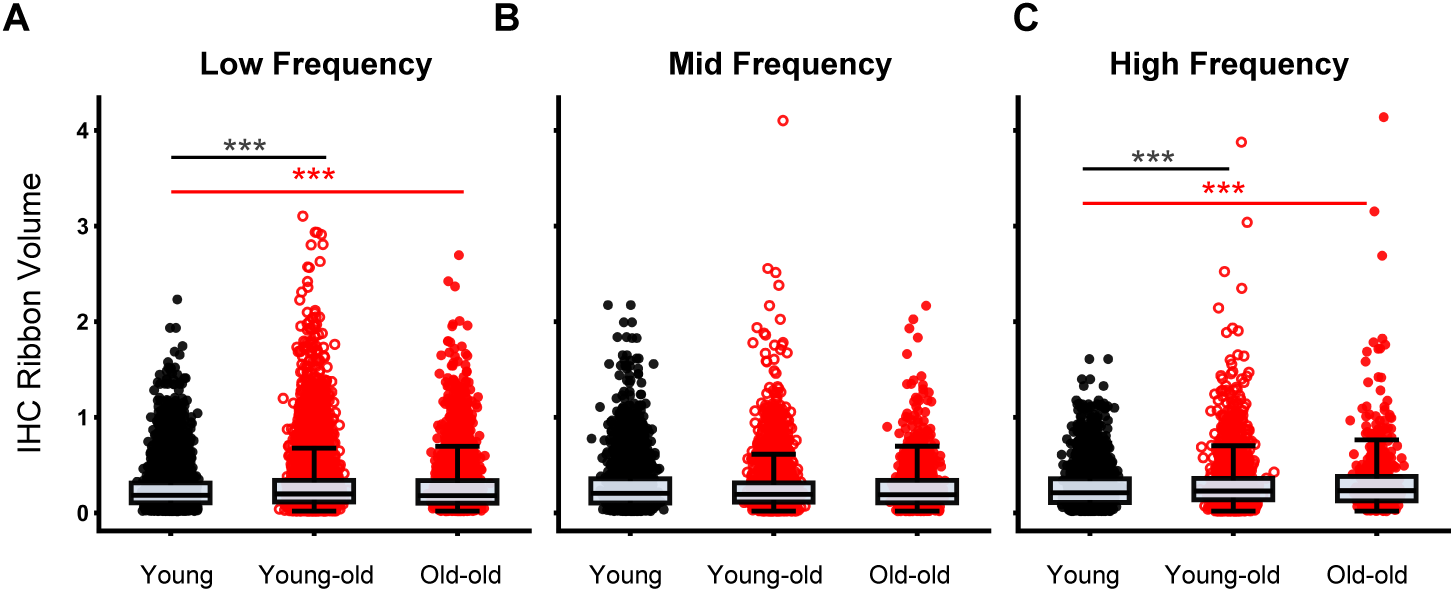
Patterns of IHC synaptic ribbon hypertrophy in aged macaques. (A-C): Swarm and box plots illustrating distribution of IHC ribbon volumes (μm^3^) for young (gray circles), young-old (red hollow circles), and old-old (red solid circles) at low (A, 0.125-2 kHz), mid (B, 4-8 kHz), and high (C, 16-32 kHz) frequencies. Significant p-values are displayed with an asterisk scheme (*: *p* < 0.05, **: *p* < 0.01, ***: *p* < 0.001).

Variability in ribbon volume also increased with age: ranges were broader in older groups, especially at high frequencies. Notably, group differences are driven primarily by the upper tail (top ≈5%) of each distribution, indicating that most ribbons retain similar volumes across ages. Although mid frequencies showed no statistically significant group differences, young-old animals trended toward hypertrophy and displayed greater high-end variability than young macaques.

### DPOAE Amplitudes Decline and Thresholds Elevate with Age

To determine whether aging impacts OHC function, we measured DP-grams using a clinically relevant stimulus condition (primary tone levels of 65/55 dB SPL and a frequency ratio of 1.22). As shown in Figure 4A, young macaques exhibited robust DP responses across frequency, while DP amplitudes declined markedly with age (all *p* < 0.001, Welch’s ANOVA and Games-Howell post-hoc). In the old-old group, DPs were absent on average across many high frequencies, suggesting significant age-related OHC dysfunction.

**Figure 4:**
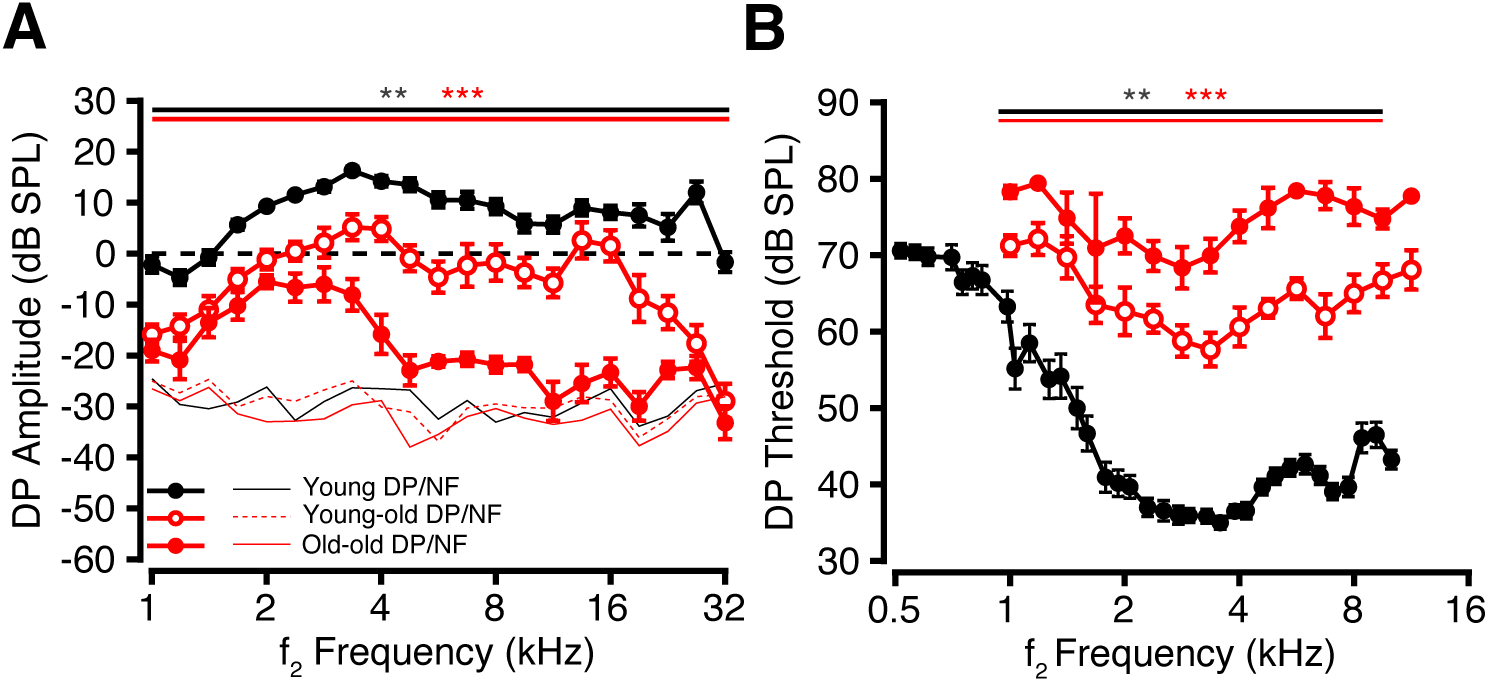
Age-related decreases in DPOAE amplitude and increases in DPOAE threshold. DPOAE amplitudes and thresholds using stimulus parameters (*f*_2_/*f*_1_ = 1.22; *L*_1_ - *L*_2_ = 10 dB). Young (black), young-old (unfilled red), and old-old (filled red) DP (Distortion Product), NF (Noise Floor), and threshold displayed. (A) Average DP amplitude (dB SPL) (±SEMs) as a function of *f*_2_ frequency at stimulus parameter, *L*_1_/*L*_2_ = 65/55 dB SPL. (B) Average *f*_2_ level at DP threshold (dB SPL) (±SEMs) as a function of *f*_2_ frequency. Significant p-values are displayed with an asterisk scheme (*: *p* < 0.05, **: *p* < 0.01, ***: *p* < 0.001).

To further examine OHC sensitivity, we measured DPs at a range of lower stimulus levels to determine DP thresholds across frequencies. Young macaques showed low DP thresholds across the frequency range, whereas thresholds were progressively elevated (*p* < 0.001 except 1.67 kHz, *p* = 0.15, Welch’s ANOVA and Games-Howell post-hoc) in older animals (Figure 4B), consistent with a loss of cochlear amplifier function and further supporting OHC impairment with age.

### Broadband ABR Responses Reveal Age-Related Processing Deficits

Auditory Brainstem Responses (ABRs) were next analyzed across various stimulus parameters in the different age cohorts (Figure 5). Representative broadband chirp waveforms show that thresholds and latencies tend to increase while amplitude decreases with age (Figure 5A). Consistent with the trend for DPs, ABR thresholds increased with age across all frequencies and broadband stimuli (Figure 5B, all *p* < 0.05, Welch’s ANOVA and Games-Howell post-hoc) and continued to increase between age cohorts from 4-16 kHz (*p* < 0.05, Welch’s ANOVA and Games-Howell post-hoc). To quantify this relationship, ABR thresholds were plotted as a function of age across multiple tone burst frequencies (Figure 5C-F). A positive linear correlation between age and threshold is seen at 4, 8, 16, and 32 kHz, where thresholds exceed the young average plus 1 SEM. These effects were statistically significant across a broad range (2-32 kHz in octave steps; data not shown for all frequencies) of tested frequencies (*p* < 0.05, OLS Regression).

**Figure 5:**
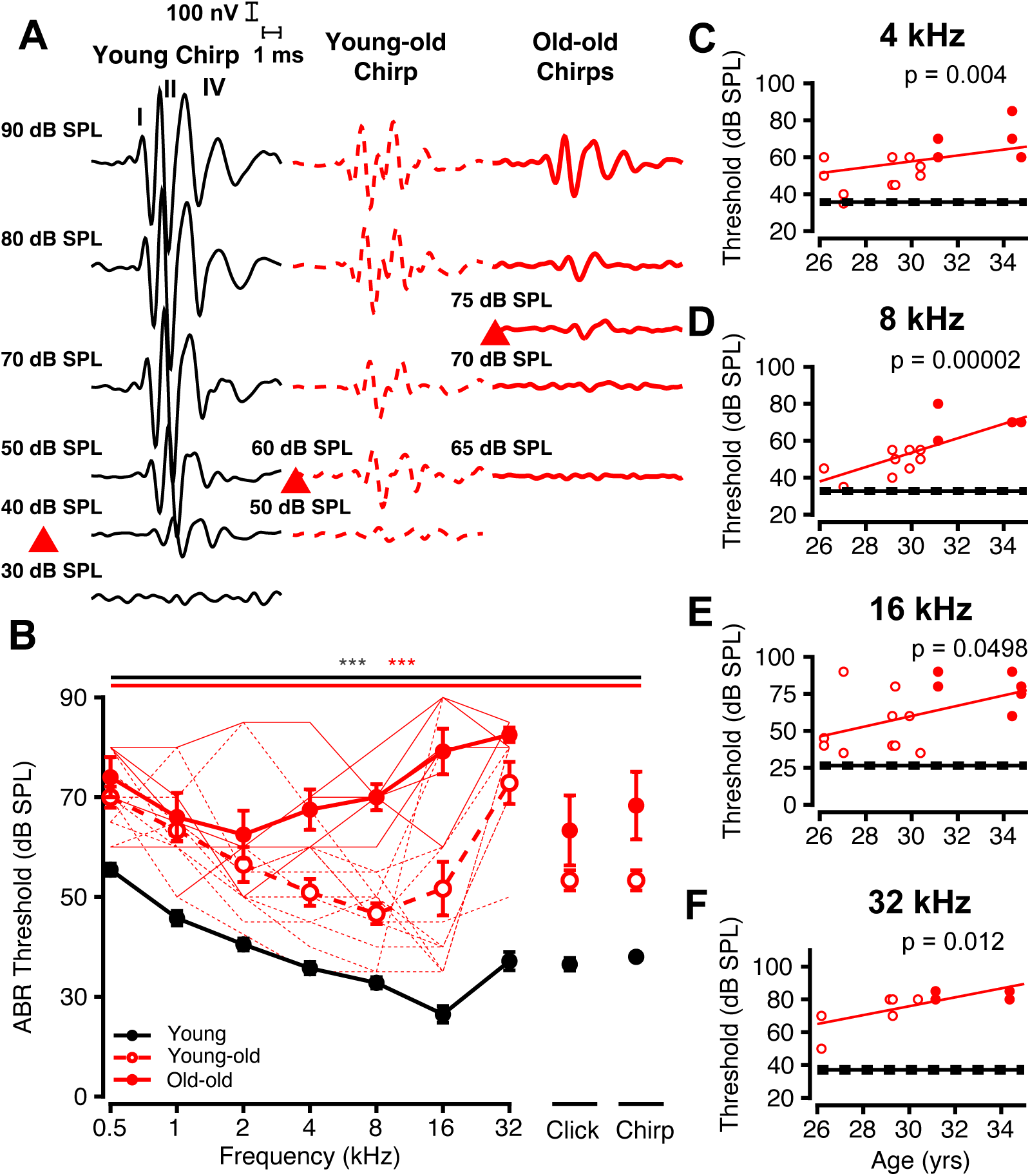
Age-Related Progressive Elevations in ABR Thresholds. ABR Sample traces, audiograms, and threshold as a function of age. (A) shows ABR sample traces of Young (black line), young-old (red dashed line), and old-old (red solid line) at different sound levels of tone burst and broadband stimuli. Threshold sample traces are noted in a red box. (B) Average ABR threshold to tone pips as a function of frequency, and thresholds to broadband clicks and chirps for young (black), young-old (unfilled red), and old-old monkeys (filled red symbols). Individual old-old (solid red) and young-old (dashed red) monkey ABR thresholds are plotted, and error bars represent one SEM. Solid (old-old) and dashed (young-old) lines are individual macaque traces for aged groups. (C-F) Threshold as a function of age, where the average young threshold is a solid black line (±SEMs, dashed black line). Young-old (unfilled red circles) and old-old (filled red circles) are plotted as points for (E) 4 kHz, (F) 8 kHz, (G) 16 kHz, and (H) 32 kHz. Solid red line represents linear fit for age vs. threshold. Significant p-values are displayed with an asterisk scheme (*: *p* < 0.05, **: *p* < 0.01, ***: *p* < 0.001).

### Chirp responses Decrease and Delay with Age

To further investigate age-related auditory deficits, we analyzed ABRs to chirp stimuli presented at varying sound levels. Macaque-specific chirps are composed of 0.5-32 kHz frequency components that match the latency differences across frequency to activate the entire cochlea simultaneously (Figure 6A).

**Figure 6:**
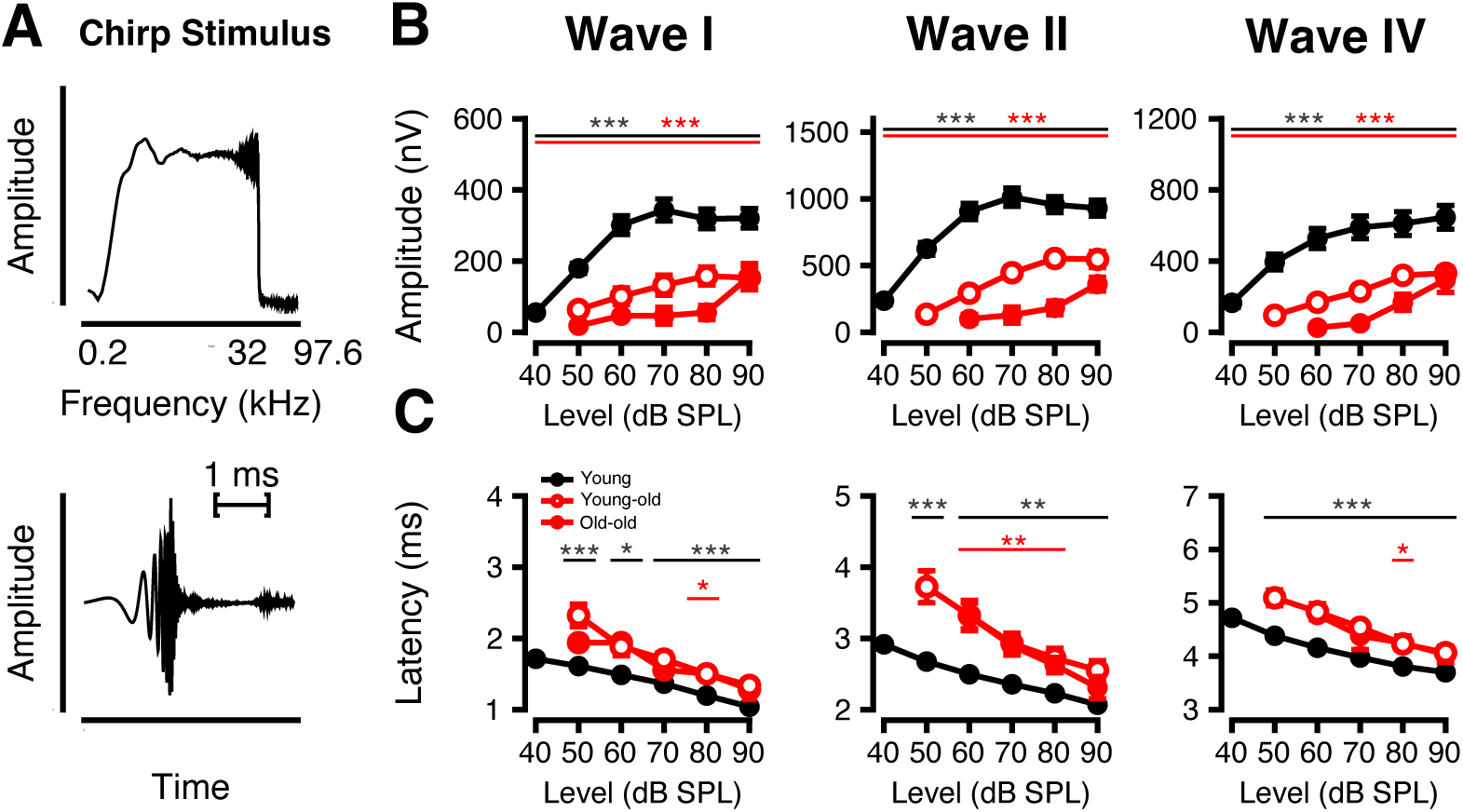
Age-related decreases in ABR amplitude and increases in ABR latency in response to chirps. ABRs in response to macaque-specific chirps as a function of stimulus sound level (dB SPL). (A) Frequency spectrum (top) and waveform (bottom) of macaque-specific chirp generated from normative macaque data. Response amplitudes (B) and latencies (C) as a function of level (±SEMs) are shown for each primary macaque waveform component (Waves I, II, and IV). Significant p-values are displayed with an asterisk scheme (*: *p* < 0.05, **: *p* < 0.01, ***: *p* < 0.001).

Compared to young monkeys, older monkeys showed significantly reduced response amplitudes (Figure 6B, *p* < 0.05 at every level, Welch’s ANOVA and Games-Howell post-hoc) and prolonged latencies (Figure 6C, *p* < 0.05, Welch’s ANOVA and Games-Howell post-hoc) across all waveform peaks. In addition, amplitudes decreased with age, particularly at Wave II (*p* < 0.05 at all levels). Wave I and IV also showed reduced amplitude with age, but only at 80- and 70-dB SPL (*p* < 0.05). Latencies did not differ significantly between young-old and old-old macaques (*p* > 0.05 at all levels). The diminished amplitude and delayed timing of chirp responses in aged animals suggest impaired recruitment of auditory nerve fibers and reduced neural synchrony.

In addition to chirp stimuli, we assessed ABRs to clicks (Figure 7). Click-evoked amplitudes (Figure 7A) for Waves I–IV decreased with age, particularly at higher sound pressure levels. In Wave I, amplitudes were significantly reduced in both young-old and old-old macaques at 80- and 70-dB SPL (*p* < 0.05, Welch’s ANOVA with Games-Howell post-hoc). Wave II showed broader declines: amplitudes were significantly lower at all levels in old-old macaques (*p* < 0.05), and in young-old macaques at every level except 90 dB SPL (*p* = 0.156, *t* = 1.47). Additionally, old-old macaques demonstrated further reductions compared to young-old macaques at 70-, 60- and 50-dB SPL (*p* < 0.05). In Wave IV, amplitude reductions were observed in both aging groups at 70- and 60-dB SPL, with a significant decrease in old-old compared to young-old macaques at 60 dB SPL (*p* < 0.05, *t* = 2.83).

**Figure 7:**
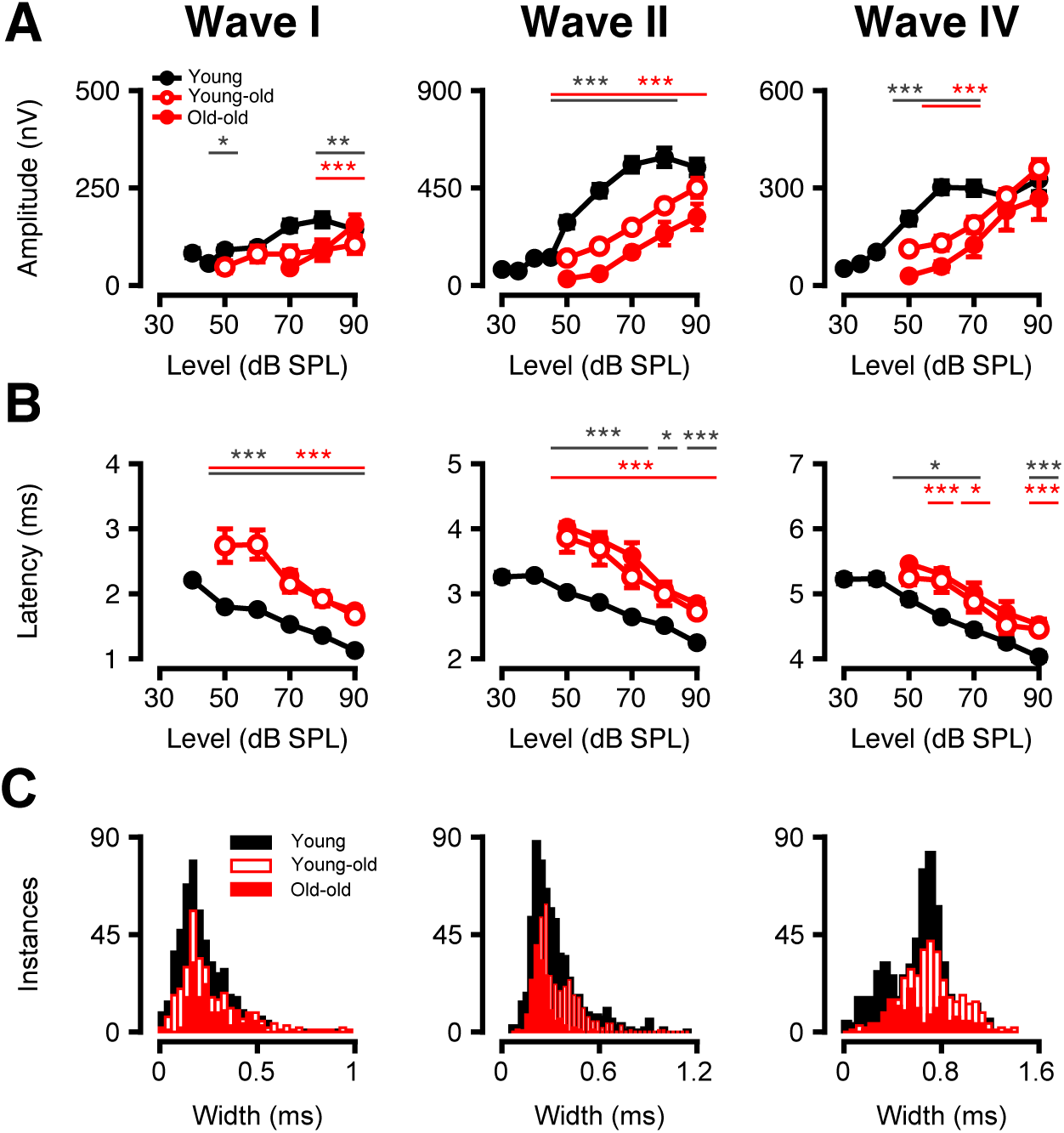
Age-related decreases in ABR amplitude and increases in ABR latency in response to clicks. Responses to clicks. ABR response amplitudes (A) and latencies (B) to macaque-specific clicks as a function of level (±SEMs) are shown for each primary macaque waveform component (I, II, IV) for clicks presented at 27.7/s. (C) Distribution of the half-maximal width of ABR Waves I-IV in response to clicks at 90 dB SPL; an intermediate group size (64 trials taken from 2048 averaged trials) was sampled from each primary wave. Significant p-values are displayed with an asterisk scheme (*: *p* < 0.05, **: *p* < 0.01, ***: *p* < 0.001).

As shown in Figure 7B, click-evoked response latencies increased with age in Waves I and II across all stimulus levels (*p* < 0.05, Welch’s ANOVA with Games-Howell post-hoc). Wave IV followed a similar trend, except at 80 dB SPL, where latencies in both young-old (*p* = 0.080, *t* = 1.87) and old-old macaques (*p* = 0.052, *t* = 2.41) were trending upwards but did not reach statistical significance. Notably, no significant latency differences were detected between young-old and old-old macaques at any level.

Waveform morphology also differed between age groups, as shown in the distribution of wave widths (Figure 7C). When analyzing the half-maximal widths of Waves I, II, and IV using small trial averages (64 trials), we found that Waves I-IV were significantly altered in older monkeys (*p* < 0.01, Kruskal-Wallis and Mann Whitney U post-hoc). Specifically, the old-old and young-old Wave I was broader than in young (right shift, *p* < 0.01), but there was no statistical difference between the two older cohorts (*p* > 0.05). Waves II and IV initially broaden in young-old macaques relative to young monkeys (*p* < 0.001), but both waves further compress in the old-old macaque cohort to values even smaller than in the young ears (*p* < 0.001). These results suggest that aging alters central auditory gain in response to peripheral delays and timing imprecision, potentially reflecting compensatory changes in neural synchrony and temporal processing across the auditory pathway.

### Clicks across Presentation Rates Differ with Age

To assess temporal processing capabilities across age, we measured ABRs to clicks presented at increasing rates ranging from 27.7 to 200 stimuli per second (Figure 8). The resulting waveforms significantly differed between age groups in their amplitudes and latencies at all presentation rates tested (Figure 8A). Compared to young monkeys, both young-old and old-old macaques exhibited a general reduction in response amplitude (Figure 8B) and an increase in peak latency (Figure 8C) across all click rates at 90 dB SPL. Young-old Wave I amplitude was significantly reduced from 125 to 200 stimuli per second (*p* < 0.05, Welch’s ANOVA with Games-Howell post-hoc), but old-old Wave I amplitude was only reduced at 166.6/s (*p* < 0.05, *t* = 2.3957). Wave I latencies showed more widespread changes, with young-old and old-old latencies significantly increasing relative to young macaques (*p* < 0.05). Wave II amplitudes were significantly reduced for all rates except for old-old macaques at 27.7/s (*p* = 0.13, *t* = 1.5788). Wave IV click amplitude did not differ significantly beyond young-old values at 27.7/s (*p* < 0.05, *t* = 3.0897). Wave II/IV latencies were less robust; while young-old latency still increased at all frequencies (*p* < 0.05), old-old latency did not significantly differ at 57.7/s and 125-200/s for Wave II and 57.7-166.6/s for Wave IV (*p* > 0.05). Young-old and old-old click rates did not significantly differ from each other (*p* > 0.05).

**Figure 8:**
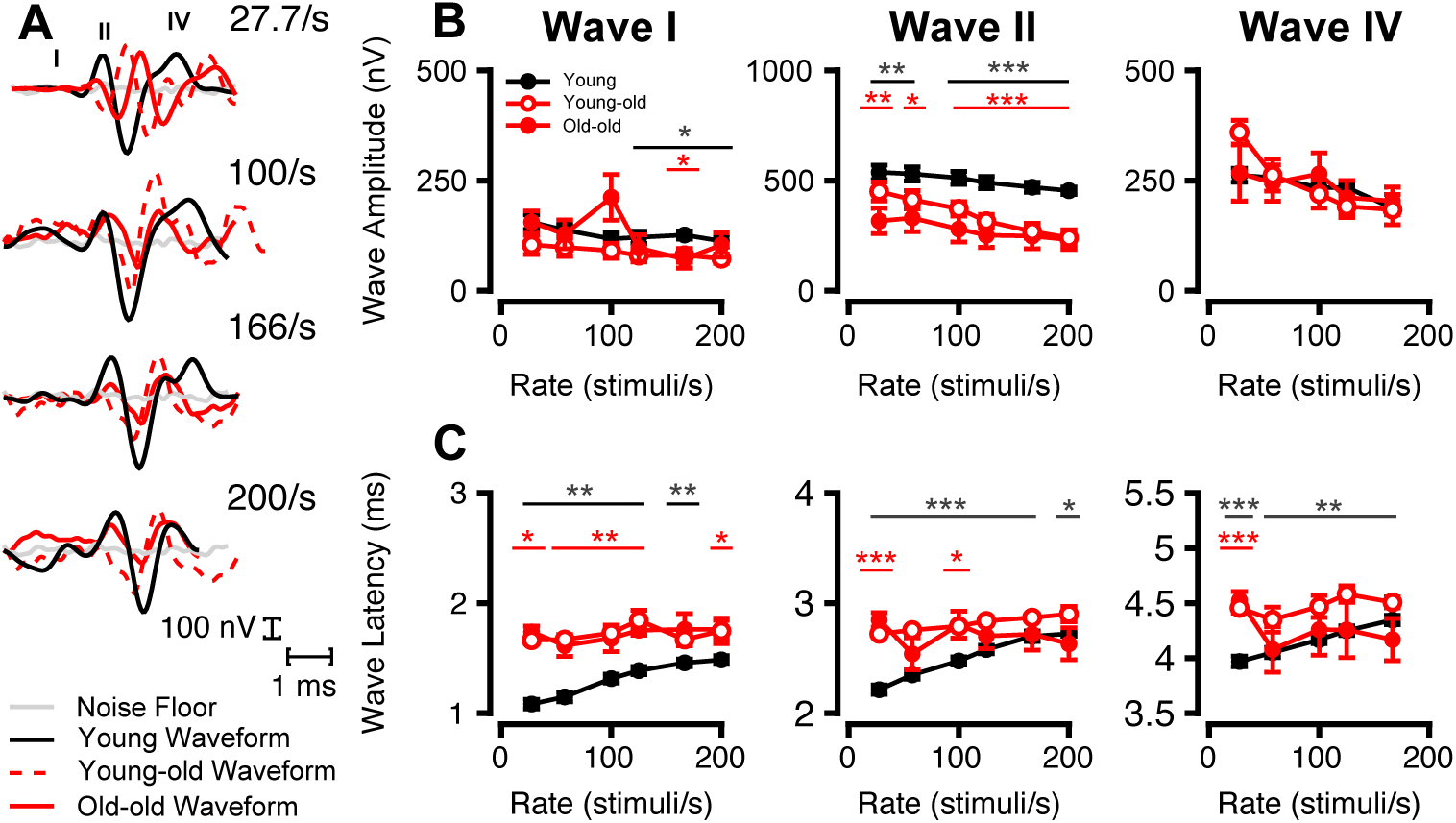
Aged Macaque Click Responses change less with Increasing Presentation Rate. Effect of click presentation rate on ABR wave amplitude and latency of different age cohorts. (A) Example traces from 1 young, 1 young-old, and 1old-old monkey in response to clicks at 90-dB SPL of increasing presentation rate (27.7/s-200/s; from top to bottom, respectively). young (solid black line), young-old (dashed red line), and old-old (solid red line) macaque cohorts are overlaid. (B) Average ABR amplitude (nV) (±SEMs) in response to 90-dB SPL clicks as a function of presentation rate (stimuli/second) for primary macaque waveform components. (C) Average ABR latency (ms) (±SEMs) in response to 90-dB SPL clicks as a function of presentation rate (stimuli/second) for primary macaque waves. Significant p-values are displayed with an asterisk scheme (*: *p* < 0.05, **: *p* < 0.01, ***: *p* < 0.001).

Ordinary Least Squares linear regression analyses incorporating an interaction term between click rate and age group revealed several significant effects (all *p* < 0.05). For Wave I and II amplitudes, the slope significantly increased in young-old relative to young macaques (*p* < 0.01), indicating enhanced rate-dependent amplitude adaptation. In contrast, Wave I and II old-old rate-dependent slopes were not significantly different from those of young (*p* > 0.05), which may be due to diminished amplitudes at lower click rates as well.

For latency responses (Waves I–IV), both aged macaque groups showed significantly flatter slopes compared to young macaques (*p* < 0.01), reflecting reduced temporal adaptation with increasing stimulus rate. These findings showcase age-related degradation in both ABR amplitude and latency dynamics, highlighting diminished neural synchronization and temporal resolution with age.

### Frequency-Specific ABR Responses Show Evidence of Recruitment

In addition to broadband analyses, we also examined ABRs to tone burst stimuli ranging from 0.5 kHz to 32 kHz (Figure 9). Unlike broadband stimuli, tone burst-evoked ABR amplitudes at 90 dB SPL did not significantly differ across age cohorts at any Wave I frequency (Figure 9A, *p* > 0.05, Welch’s ANOVA and Games-Howell post-hoc). Any differences may be masked by the high inter-ear and inter-subject variability in macaque tone-burst ABRs (Stahl et al., 2022). Notably, while young-old macaques exhibited a reduction in Wave II and Wave IV amplitudes (0.5-8 kHz, *p* < 0.001), this decrease was not sustained in the old-old cohort, which appeared to partially recover response strength (*p* > 0.05) in these frequencies and reduce amplitude in ranges that were previously not significantly different from young or young-old macaques (Wave II 16 kHz, *p* < 0.05, *t* = 4.4629; Wave IV 16-32 kHz, *p* < 0.01).

**Figure 9:**
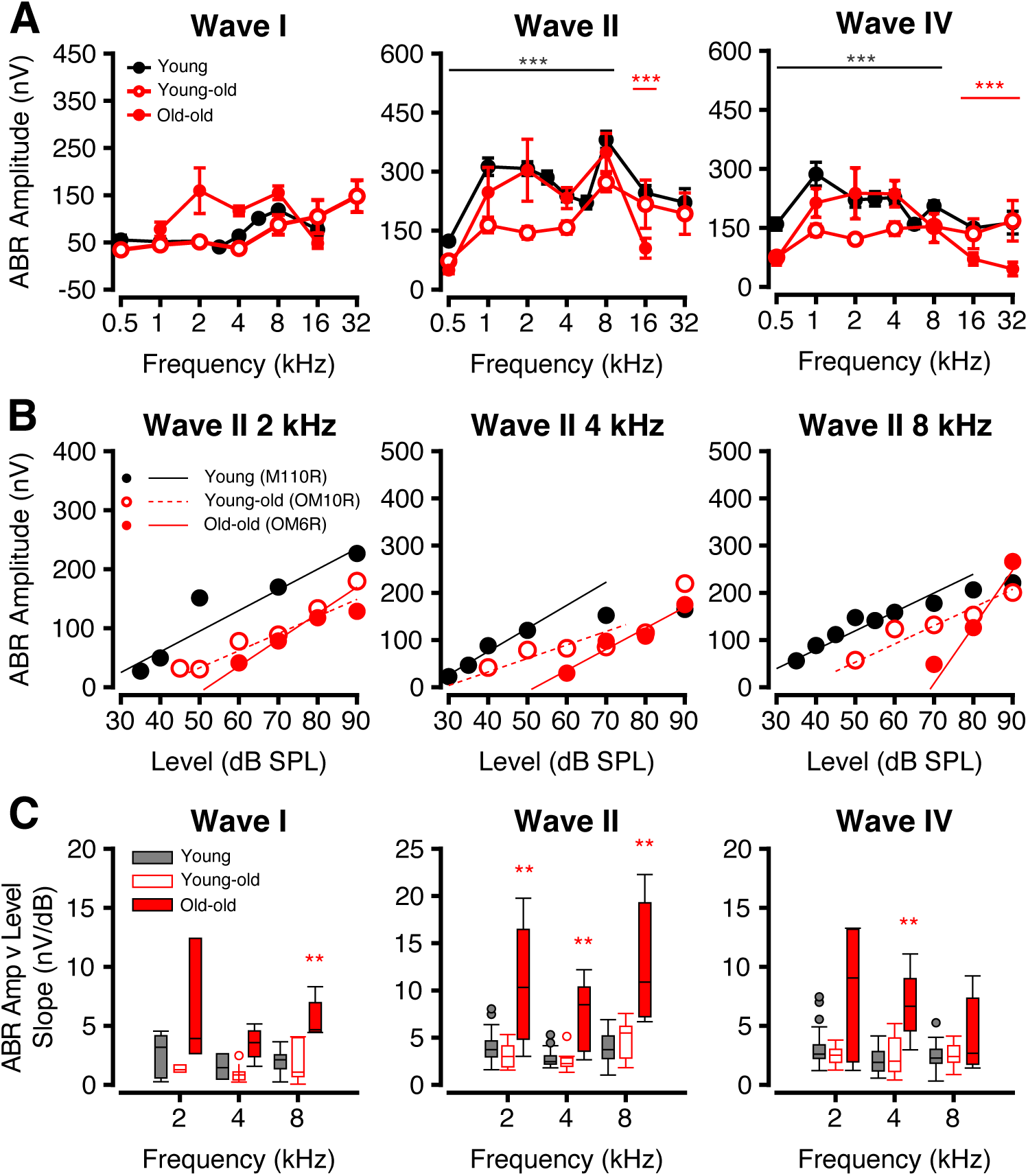
Tone-specific responses show paradoxical stability in old monkeys. ABR tone burst responses across various frequencies (kHz) to stimuli at different sound levels. (A) Average ABR amplitude (nV) (±SEMs) at 90 dB SPL at frequencies 0.5-32 kHz for each waveform component for young (filled black), young-old (unfilled red), and old-old (filled red) age cohorts. (B) Representative scatter plots of Wave II ABR amplitude (nV) as a function of sound level of young (black circles, black solid line), young-old (red unfilled circles, red dashed line), and old-old (red filled circles, red solid line) macaques at 2, 4, and 8 kHz. (C) Boxplots of linear slope (Amplitude as a function of level) distributions of young (dark grey), young-old (unfilled red), and old-old (filled red) for ABR Waves I-IV. Significant p-values are displayed with an asterisk scheme (*: *p* < 0.05, **: *p* < 0.01, ***: *p* < 0.001).

To investigate the sensitivity of the auditory system across sound intensities, we plotted ABR Wave II amplitude, the most robust ABR wave in macaques (Stahl et al., 2022), as a function of stimulus level for 2, 4, and 8 kHz tones. This analysis revealed distinct differences in slope between age groups (Figure 9B). Young-old monkeys exhibited reduced amplitudes (*p* < 0.05, Welch’s ANOVA and Games-Howell post-hoc) without significant slope changes (*p* > 0.05, Kruskal-Wallis and Mann Whitney U post-hoc) across all waves, suggesting diminished dynamic range and impaired encoding of sound intensity: hallmarks of a degraded auditory system with limited capacity to discriminate between sound levels.

Slope distributions for each age group by wave and frequency were averaged as shown in Figure 9C. Relative to young and young-old macaques, which maintained relatively similar slopes, old-old monkeys showed steeper amplitude-intensity slopes at Wave I 8 kHz, all Wave II frequencies, and Wave IV 4 kHz (*p* < 0.01, Kruskal-Wallis and Mann-Whitney U post-hoc).

### Physiology and Histology deficits correlate

Figure 10B plots the ABR threshold shifts (re young macaques from Stahl et al. 2022) against OHC. All three rows show significant negative exponential correlations with shift, i.e. as ABR threshold shifts increase, OHC survival decreases (Exponential correlation, all *p* < 0.0001). A linear mixed effects model revealed that, even though all OHC rows correlate with ABR shifts individually, OHC row 3 (furthest from IHCs) carries the strongest independent signal (partial F = 8.41, *p* = 0.0049) with OHC row 1 also marginally contributing (partial F = 3.39, *p* = 0.069).

**Figure 10:**
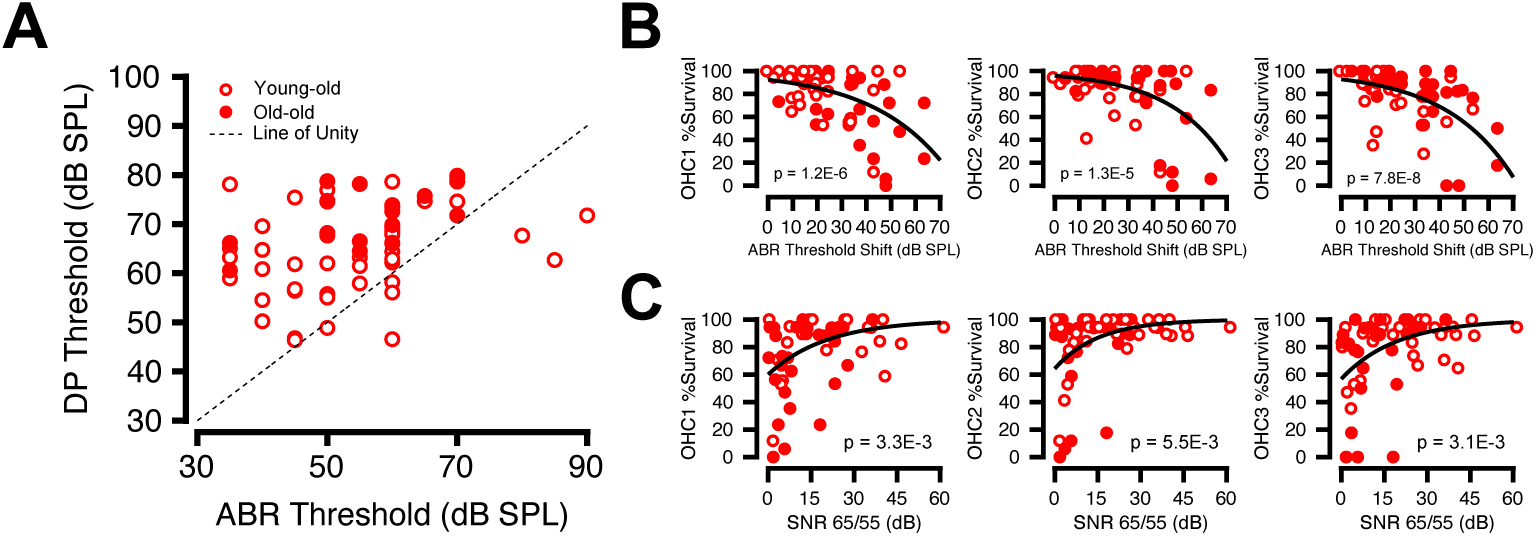
Correlation between Structural Loss and Physiological Impairment. Scatter plots correlating histological and physiological metrics of young-old (unfilled red circles) and old-old (filled red circles) macaques. (A) Correlating DP to ABR; dotted black line represents line of unity (y = x). (B) OHC Rows 1-3 % (*r*_1_ = −0.5207; *r*_2_ = −0.4750; *r*_3_ = −0.5666) for OHC frequency-matched ABR threshold shift. (C) DP amplitude (*L_1_/L_2_* = 65/55 dB SPL, *f_2_/f_1_* =1.22) as a function of OHC % survival by row. Solid black line represents linear fit (*r*_1_ = 0.3674; *r*_2_ = 0.3488, *r*_3_ = 0.3697) for DP Amplitude at 65/55 dB SPL vs. OHC Survival % by row. All *p* values listed on the graph represent model fit. Significant p-values are displayed with an asterisk scheme.

Figure 10C shows DP SNR (DP amplitude – NF) at each *f_2_* frequency plotted against OHC row. As for ABR threshold shifts, the DP SNR correlates with OHC survival. OHC survival was higher in all 3 rows when the DP SNR was higher. (mean r = 0.3620, all *p* < 0.01).

Here, a linear mixed effects model revealed that no single row dominated the relationship, indicating a global loss in OHCs led to decreased SNR amplitude.

### Structural and Functional Auditory Decline Associated with Early Cognitive Impairments

Given the comprehensive auditory profiling conducted in this study, we next asked whether peripheral auditory deficits were related to cognitive performance through a delayed-match-to-sample (DMS) task on the same animals – a unique opportunity made possible with a macaque model (Figure 11). Notably, the DMS task was entirely visual, allowing correlations to emerge without auditory task demands. The DMS task delay was adaptively adjusted each trial so that performance converged onto a 79% level (Pennington et al., 2025). We call this delay the “memory threshold delay” which quantifies the monkeys’ overall performance of the task. To further isolate different cognitive functions contributing to task performance, we modeled each monkey’s per-trial responses across delays using a mixed-effect logistic regression model in which delay was treated as a categorical random-effect factor (see Methods). We identified a delay-averaged “reliability” factor controlling how sensitive the monkey’s choice was to the target layout, in addition to a left-right “bias” factor independent of target layout. Reliability captures sensory and memory fidelity while bias captures lateralized attentional or motor deficits, both measured in log-odds.

**Figure 11:**
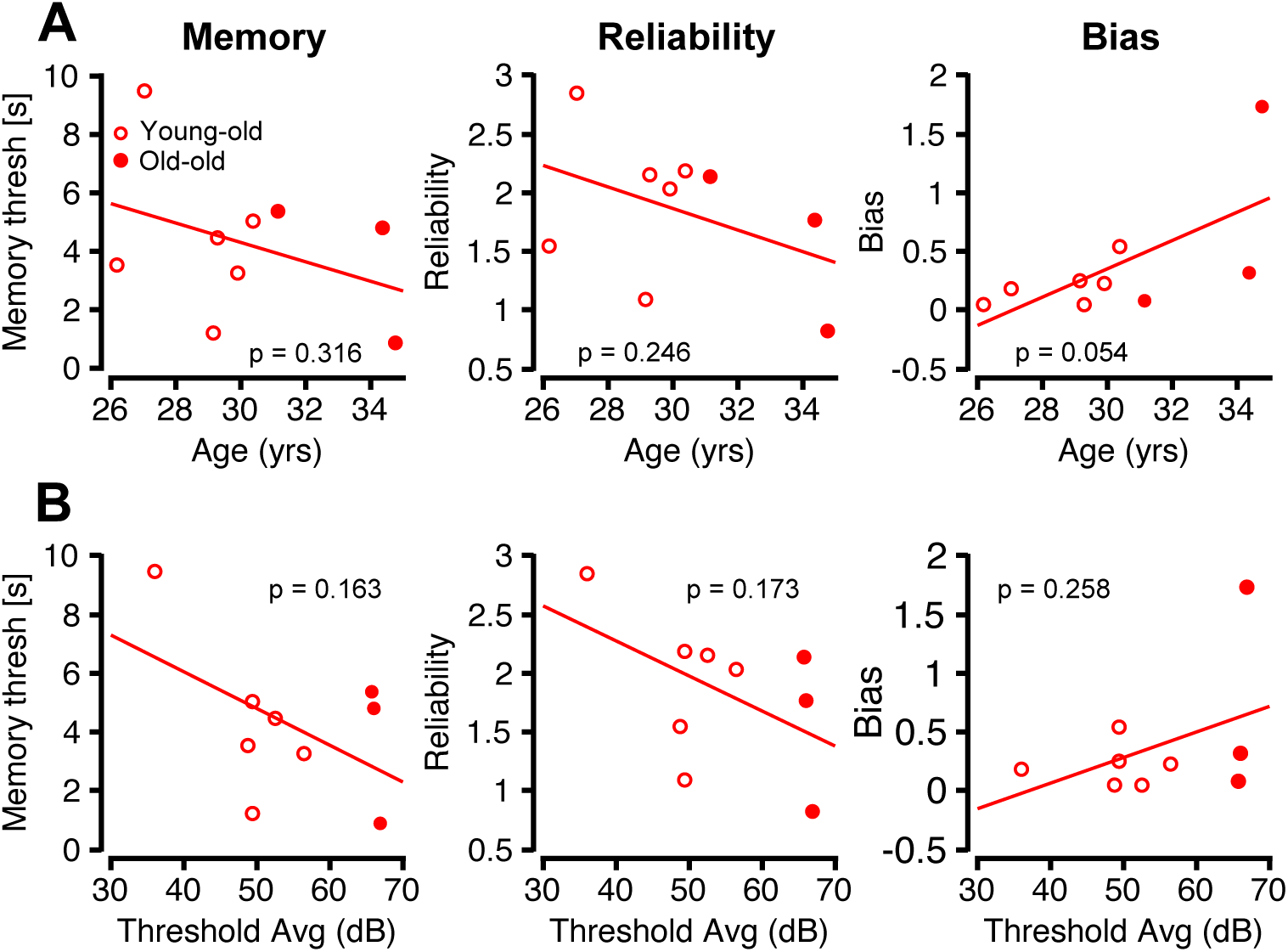
Cognition declines with age and auditory function. Old macaque cognitive metrics from a delayed match-to-sample (DMS) task are plotted as a function of age and threshold, *p*-values displayed in each panel. (A) Old macaque cognitive metrics plotted as a function of age. (B) Each cognitive metric plotted against the average threshold of the ABR at 4, 8, and 16 kHz.

Figure 11A establishes a relationship between cognitive metrics and age. The data suggests that older monkeys tended to have lower memory threshold delay and reduced reliability, alongside a subtle increase in bias (*p* > 0.05, OLS regression). These trends suggest aging may lead to less stable and efficient cognitive processing.

Figure 11B extends the trends in cognitive metrics with measures of auditory integrity/perception indexed with average ABR tone burst thresholds at 4, 8, and 16 kHz, the three most sensitive frequencies (Stahl et al. 2022). Macaques with higher thresholds tended to have lower memory thresholds, reduced reliability, and increased bias in the DMS task (Figure 11C left, middle and right, respectively). Individual tone burst thresholds, along with other combinations of averaged tone burst thresholds, were also compared with each cognitive metric, but no trending relationship was found (not shown). Though these relationships did not reach statistical significance (*p* > 0.05, OLS regression), the directionality of all three trends supports the idea that auditory decline may co-occur with, and potentially contribute to, age-related changes in cognition.

## Discussion

This study investigated age-related structural and physiological changes in the auditory system of rhesus macaques to identify early biological markers of hearing decline and their relevance to cognitive dysfunction. Using immunohistochemistry, DPs, ABRs and delayed matching-to-sample tasks, we found evidence that aging is associated with progressive cochlear degeneration, worsened auditory processing, and signs of cognitive decline. These alterations mirror the hearing and cognitive challenges often observed in older human populations (Gates & Mills, 2005). Importantly, because these monkeys were housed in a quiet laboratory environment (Mcleod et al., 2022), this model isolates aging-related effects from noise-induced hearing loss, a confounding factor in human aging studies (Valderrama et al., 2018). With their relatively long lifespan, phylogenetic similarity to humans, and trainability on behavioral tasks, rhesus macaques provide a powerful framework for studying sensory and cognitive aging.

Histological analysis revealed a progressive degeneration of both OHCs and IHCs with increasing age, consistent with prior observations in macaques (Engle et al., 2013). OHC loss was overall more extensive than IHC loss, mirroring trends seen in rodent and human studies (Spongr et al., 1997; Wu et al., 2019). However, significant IHC loss was also observed in basal cochlear regions, indicating that while OHCs are preferentially affected, IHC degeneration also contributes to auditory decline in advanced age. Although our data showed significant correlation between OHC losses and ABR threshold elevations, the best fit was exponential in nature (Figure 10), where only the largest threshold shifts were associated with hair cell loss. This contrasts with data from aging humans where the relation between OHC losses and audiometric threshold was well fit by a straight line, i.e. where even modest threshold shifts were on average associated with significant OHC loss (Wu et al., 2020).

Distortion products were also attenuated in the aging macaques. Indeed, they were close the noise floor at mid to high frequencies in the oldest animals, despite no corresponding reduction in OHC number within the same cochlear regions. In some animal and human studies of noise-induced or age-related hearing loss, the threshold shifts are highly correlated with damage to the stereocilia (Liberman and Dodds 1984; Wu and Liberman 2024). In the present study, some of the old-old ears showed damaged stereocilia, however there remained many ears where the stereocilia appeared normal. Pronounced threshold elevations have been observed in other studies in the absence of proportional OHC loss (Stebbins et al., 1979; Moody et al., 1978; Hawkins et al., 1976; Xia et al., 2022). The age-related decline in DPOAEs and ABRs may reflect dysfunction of surviving OHCs, such as impaired electromotility.

Additional non-sensory factors such as strial dysfunction may also contribute to the early functional decline in the macaque.The threshold shifts in our old monkeys (Fig. 4B) showed a relatively flat configuration, with elevated low-frequency thresholds and mild high-frequency roll-off—an audiometric pattern that has been suggested, based on work in aging gerbils, to indicate damage to the stria vascularis and reductions in the endolymphatic potential (Vaden et al 2022; Lang et al,, 2023). Recent histopathological work in human temporal bones has confirmed that low-frequency threshold elevation is an indicator of strial damage (Kaur et al., 2023).

Prior work with noise-exposed cats suggested that first row OHCs were more vulnerable to stereocilia damage and made larger contributions to cochlear amplification than the other rows (Liberman and Dodds, 1984), while primate noise-exposure studies report greater structural vulnerability in row 3 (Moody et al., 1978; Hawkins et al., 1976). Our findings (Figure 10B/C) suggest a comparatively stronger contribution to thresholds from row 3. These results collectively reinforce the concept that cochlear dysfunction reflects a progression from early functional impairment in surviving OHCs to later structural degeneration, with row-specific contributions that may vary across species.

Synaptic ribbon counts also showed a relatively modest age-related decline in the macaque model. This contrasts with human data, where cochlear neural degeneration (measured as loss of auditory nerve peripheral axons) declined steadily with age, with an average 40% loss across the entire audiometric frequency range by age 70 (Wu et al.2019). Comparable findings have been observed in mouse models, where there is a steady age-related reduction in synapse counts across the entire cochlea, with an average loss of 50% by the end of life (Sergeyenko et al 2013). In both mice and humans, these synaptic losses were accompanied by larger OHC losses than seen in the present study. Prior macaque work shows that there can be massive synaptopathy after noise exposures that destroy large numbers of OHCs (Valero et al., 2017). If the macaques in the present study really were near the end of their lifespan, it suggests that their cochleae are less vulnerable to aging than either mice or men, although we did see relative increases in ribbon volume from base to apex, analogous to that reported in aging rodents and humans (Jeng et al., 2020; Sergeyenko et al., 2013).

ABR suprathreshold responses provided further evidence of age-related peripheral and central auditory decline. As a noninvasive assay of neural synchrony and conduction (Møller & Jannetta, 1985), ABRs showed reduced amplitudes, and prolonged latencies in waves I, II, and IV with age, as seen in human populations (Konrad-Martin et al., 2012). As described by Ng et al. (2015), ABR amplitudes are a function of the number and synchrony of neural units firing. ABR latency reflects the neural transmission time of auditory signals. The age-related amplitude reductions likely indicate loss or desynchronization of auditory nerve fibers, whereas prolonged latencies and wave dispersion suggest impaired temporal precision and slower conduction through central auditory pathways including the cochlear nucleus, superior olivary complex, and inferior colliculus.

Macaque-specific chirp stimuli are designed to activate the entire cochlea at the same time by presenting a frequency sweep with delays derived to compensate for the traveling wave of excitation along the cochlear spiral (Wegner & Dau, 2002). In the old-old group, chirp waves I–IV showed latencies relative to young macaques, suggesting degraded temporal fidelity in early brainstem and midbrain relays (Lasky et al., 1995; Gray et al., 2014). These findings align with electrophysiological studies in aging rodents that demonstrate cellular-level changes, such as diminished inhibition, increased spontaneous activity, and impaired temporal coding in these same regions (Stebbins et al., 2016; Kersbergen et al., 2023; Khouri et al., 2011; Parthasarathy et al., 2010).

Click-evoked ABRs corroborated chirp findings, showing pronounced declines in Waves I-IV as seen in other aged macaque studies (Ng et al., 2015). This may be suggestive of age-related desynchronization of auditory nerve fibers firing (Sergeyenko et al., 2013) that may influence function of higher order structures. Older animals also exhibited steeper tone-specific amplitude-intensity slopes, which may reflect enhanced neural recruitment at suprathreshold levels – a possible compensatory mechanism for temporal discrimination loss (Marchetta et al., 2020).

These auditory declines may also serve as early indicators of broader neurocognitive vulnerability. Epidemiological studies consistently link age-related hearing loss with elevated risks of cognitive impairment and dementia (Loughrey et al., 2018; Guo et al., 2023), with even mild hearing impairment being significantly associated with cognitive decline (Conceição Santos de Oliveira et al., 2023). Our observation that aged macaques with greater auditory dysfunction also performed more poorly on DMS tasks supports the idea that sensory degradation can precede and potentially contribute to cognitive decline. Given the anatomical and behavioral similarities between rhesus macaques and humans (Chiou et al., 2020), these results provide a foundation for more detailed studies of age-related auditory and cognitive decline in nonhuman primates.

With its extended lifespan, human-like auditory range, and capacity for multimodal assessment, the rhesus macaque offers a powerful platform for evaluating interventions and probing mechanisms across human-relevant aging trajectories. Ultimately, these findings not only refine our understanding of auditory aging, but they also position the rhesus macaque as a pivotal model for testing therapeutic strategies aimed at preserving hearing and cognition in the aging brain.

## ACKNOWLEDGEMENTS

The authors are especially grateful to Mary Feurtado for her expertise and assistance with anesthesia maintenance in a terminal experiment. This work was supported by NIH–NIDCD R01 DC015988 (MPI: R. Ramachandran and B. Shinn-Cunningham) and RF1 AG 060754 (PI: Christos Constantinidis).

The authors have no conflicts of interest, financial or otherwise, to disclose.

